# Modular multiwell viscoelastic hydrogel platform for two- and three-dimensional cell culture applications

**DOI:** 10.1101/2023.10.09.561449

**Authors:** Mackenzie L. Skelton, James L. Gentry, Leilani R. Astrab, Joshua A. Goedert, E. Brynn Earl, Emily L. Pham, Tanvi Bhat, Steven R. Caliari

**Affiliations:** Department of Biomedical Engineering, University of Virginia, Charlottesville, Virginia 22903; Department of Psychology, University of Virginia, Charlottesville, Virginia 22903; Department of Chemical Engineering, University of Virginia, Charlottesville, Virginia 22903

**Keywords:** hydrogels, multiwell plate, high-throughput, disease modeling

## Abstract

Hydrogels have gained significant popularity as model platforms to study the reciprocal interactions between cells and their microenvironment. While hydrogel tools to probe many characteristics of the extracellular space have been developed, fabrication approaches remain challenging and time-consuming, limiting multiplexing or widespread adoption. Thus, we have developed a modular fabrication approach to generate distinct hydrogel microenvironments within 96-well plates for increased throughput of fabrication as well as integration with existing high-throughput assay technologies. This approach enables *in situ* hydrogel mechanical characterization and was used to generate both elastic and viscoelastic hydrogels across a range of stiffnesses. Additionally, this fabrication method enabled a 3-fold reduction in polymer and up to an 8-fold reduction in fabrication time required per hydrogel replicate. The feasibility of this platform for cell culture applications was demonstrated by measuring both population-level and single cell-level metrics via microplate reader and high-content imaging. Finally, the 96-well hydrogel array was utilized for 3D cell culture, demonstrating the ability to support high cell viability. Together, this work demonstrates a versatile and easily adoptable fabrication approach that can support the ever-expanding tool kit of hydrogel technologies for cell culture applications.

## 1. Introduction

Hydrogels are useful tools for modeling extracellular environments and probing cellular behavior *in vitro* due to their ability to mimic native tissue properties. The direct, orthogonal control with *in vitro* hydrogel models over aspects of the extracellular matrix (ECM) that are generally coupled *in vivo* enables investigation of the individual and combined effects of various extracellular properties on cell behavior. The ability to tune key properties of the extracellular environment such as stiffness^1–3^, viscoelasticity^4,5^, cell adhesion motifs^6^, and presentation of biochemical cues^7,8^ has been demonstrated by various groups. For example, control over viscoelasticity has been shown to dictate stem cell differentiation, with faster relaxing hydrogels promoting preferential osteogenesis^4^. Yet, a primary limitation of hydrogel models is their low fabrication throughput, as these platforms are often formed individually on coverslips and cultured within large well plates or dishes. This limits the experimental space that can be explored, demanding large investments of time, hydrogel material, and reagents when seeking to conduct large-scale experiments. Thus, the development of methods to increase the fabrication throughput of hydrogel cell culture platforms is essential for widespread adoption of hydrogel platforms.

Various methods of hydrogel fabrication with increased throughput have been developed with unique sets of advantages and disadvantages centered around factors such as fabrication time and complexity^9^. Platforms such as microarrays and microwells offer adequate throughput required for large scale drug screens, but often with little to no biomimetic complexity. Conversely, gradients, microgels, and dynamic culture systems offer aspects of biological complexity that are advantageous for specific use-cases and modeling organ-level movement. However, these systems each require unique fabrication and cell culture apparatuses for implementation. One widely accessible approach is the formation of hydrogels in 96-well plates. Unlike many other high-throughput hydrogel platforms, utilizing hydrogels within 96-well plates maintains similar fabrication methods and culture conditions to traditional hydrogel cell culture models, making them easily adapted for both current and new hydrogel users. Further, fabricating 96-well hydrogel arrays is compatible with commercially available technologies such as bottomless 96-well plates, multichannel pipettes, microplate readers, and high-content imaging systems. Thus, this approach offers a powerful yet accessible option for generating high-throughput, biologically mimetic cell culture platforms.

Considering these advantages, a few groups have developed methods for generating unique hydrogel systems within 96-well plates. Lei et al. varied ultraviolet (UV) light intensity to create a 96-well methacrylated hyaluronic acid (HA) hydrogel array to investigate the synergistic effects of stiffness and cell adhesion ligand concentration on mesenchymal stem cell behavior^10^. While this method addressed many of the described design criteria, there is a need to expand the investigative capacities of these 96-well-based hydrogels beyond two variables. Brooks et al. utilized automated liquid robotics to generate polyethylene-glycol (PEG)-based hydrogels for both two-dimensional (2D) and 3D cell culture applications^11^, and Mih et al. utilized stainless steel replicators affixed with glass squares to cast 2D polyacrylamide hydrogels in 96- and 384-well plates^12^. While these systems are powerful in scalability, there is a need for the introduction of new iterations on these fabrication techniques that increase accessibility and cater to unique hydrogel chemistries and design criteria. Specifically, there is an opportunity for a fabrication approach that is adaptable toward a wide range of hydrogel chemistries, culture dimensionality, biological questions, and subject-matter expertise.

Here, we developed a novel approach to the generation of a 96-well hyaluronic acid-based hydrogel array. Through the use of versatile thiol-ene photochemistry, we demonstrate that these hydrogels can be tuned to modulate stiffness and viscoelasticity in both 2D and 3D cell culture. This modular fabrication approach is adaptable to a range of hydrogel mechanics and components such as matrix metalloproteinase (MMP)-degradable crosslinkers and bioactive moieties, while maintaining accessibility to many new users. Finally, these culture platforms are compatible with high-throughput measurement techniques such as microplate readers and high-content imaging systems. Overall, we propose that this platform is a useful tool that satisfies a wide range of hydrogel-based cell culture needs.

## 2. Materials and Methods

### 2.1 NorHA synthesis

As previously described, HA was functionalized with norbornene groups by amine coupling of 5-norbornene-2-methylamine to a tert-butyl ammonium salt of HA (HA-TBA) with benzotriazole-1-yloxytris-(dimethylamino)phosphonium hexafluorophosphate (BOP)^13,14^. Sodium hyaluronate (Lifecore, 82 kDa) was dissolved in deionized (DI) water and reacted with Dowex 50W proton-exchange resin at 25°C for 2 h. The resin was then vacuum filtered, and the remaining solution was titrated to pH of 7.05 before being frozen, lyophilized, and evaluated using ^1^H NMR to confirm the successful production of HA-TBA (**Figure S1**). The resultant HA-TBA was then reacted with 5-norbornene-2-methylamine and BOP in anhydrous dimethylsulfoxide (DMSO) for 2 h at 25°C under nitrogen. The reaction was quenched using cold DI water and subsequently dialyzed in 6-8 kDa tubing for 3 days in salt water and 2 days in DI water. The solution was vacuum filtered to remove side products from the BOP coupling and dialyzed again in DI water for 5 days before being frozen and lyophilized for long-term storage at -20°C. The final product was analyzed through ^1^H NMR and the degree of modification was determined to be 16% (**Figure S2**).

### 2.2 CD-HA synthesis

β-cyclodextrin-modified hyaluronic acid (CD-HA) was synthesized as previously described^14,15^. First, β-cyclodextrin hexamethylene diamine (CD-HDA) was synthesized by adding dropwise p-Toluenesulfonyl chloride (TosCl) dissolved in acetonitrile to a suspension of CD in ultra-pure water (1.25:1 molar ratio of TosCl to CD) over ice. After stirring for 2 h, cold NaOH was added dropwise (3.1:1 molar ratio of NaOH to CD) to the solution before allowing the reaction to continue for an additional 30 min off ice. Ammonium chloride was added until a pH of 8.5 was reached, after which the solution was cooled on ice and centrifuged to recover the precipitate. Decanting and reprecipitation with cold water were repeated 3 times. Finally, the product was washed 3 times in cold acetone and once with cold diethyl ether and left to dry overnight. The CD-Tos product was charged with HDA (4g/g CD-Tos) and dimethylformamide (DMF) (5mL/g CD-Tos) and left to react for 12 h at 80°C under nitrogen. The final CD-HDA product was precipitated with cold acetone, washed with cold diethyl ether, and vacuum dried. The degree of modification was determined to be 73% by ^1^H NMR (**Figure S3**).

To achieve the final CD-HA polymer, CD-HDA was reacted with HA-TBA and BOP in anhydrous DMSO at 25°C for 3 h. This reaction was quenched with cold water, dialyzed for 5 days, filtered, and dialyzed for 5 more days. The purified solution was frozen and lyophilized. The degree of modification of HA was determined by ^1^H NMR to be 25% (**Figure S4**).

### 2.3 96-well plate fabrication

Glass pieces with the same thickness as a number 1.5 coverslip (0.17 mm, Schott D263) were cut to the dimensions of a 96-well plate (110 x 75 mm, S.I. Howard Glass, Worcester, MA). Bottomless 96-well plates without adhesive (Greiner Bio-One, Kremsmunster, Austria) were bound to microfluidic-grade double-sided adhesive (ARcare 90196NB, Adhesive Research, Glen Rock, PA) that was laser cut to the plate dimensions (**Figure S5**). When adhesive was applied to the bottomless 96-well plates and subsequently the glass bottom, special care was taken to ensure complete adhesion through application of gentle pressure evenly across the plate. For 2D cell culture, silicone spacer sheets of 0.5 mm thickness (Grace BioLabs, Seattle, WA) were laser cut to the 96-well geometry with a diameter of 0.5 mm less than the wells to enable some margin for improper alignment (**Figure S6**). For 3D culture, a 2 mm thick polydimethylsiloxane (PDMS) sheet of otherwise same geometry was fabricated. An alignment piece was 3D printed to assist with alignment of adhesive to plates, silicone sheet to glass, and bottomless plate to glass (**Figure S7**).

### 2.4 PDMS space fabrication

The PDMS was molded using a custom 3D-printed part (**Figure S8**). The mold was printed on an Ender-3 S1 3D printer using a 1.75 mm polylactic acid (PLA) filament (SUNLU, Guangdong, China) at a resolution of 0.2 mm layer height and 30% infill. PDMS was prepared by mixing base and curing agent at a weight ratio of 10:1. The mixture was stirred vigorously and then degassed under vacuum for 1 h. The PDMS mixture was then slowly poured into the mold to avoid the introduction of air bubbles. A clear plastic sheet was placed on top of the uncured PDMS and flattened to be flush with the mold to ensure a flat surface. The mold was then placed in an oven at 35°C for 2 h or until the PDMS was fully cured. After carefully removing the PDMS from the mold, a 6 mm biopsy punch was used to cut wells out of the PDMS at the site of the mold indentations, and the sheet was cleaned thoroughly before using for 3D hydrogel fabrication.

### 2.5 HA hydrogel fabrication

To provide a surface for the gels to adhere to, glass pieces were functionalized with free thiol groups by conjugating (3-mercaptopropyl)trimethoxysilane onto glass at 100°C following plasma treatment. Functionalized glass pieces were washed sequentially with dichloromethane, 70% ethanol, and ultrapure water to remove unreacted silane.

Hydrogels were formed through ultraviolet (UV)-light mediated thiol-ene addition similarly to previously established methods^6^. Hydrogels with elastic moduli (E’) of 1 kPa (2 wt% NorHA), 6 kPa, and 25 kPa (4 wt% NorHA) were crosslinked using dithiothreitol (DTT) at a thiol:norbornene ratio of 0.1, 0.15, and 0.4 respectively. Viscoelastic behavior was introduced into this hydrogel system by incorporating physical interactions between CD-HA and a thiolated adamantane peptide (GCKKK-Adamantane, Genscript). Viscoelastic hydrogels with E’ of 1 kPa and tan delta > 0.1 (4 wt% CDHA-NorHA) were crosslinked using DTT at a thiol:norbornene ratio of 0.05 and a CD-HA-adamantane peptide mixture at a CD:adamantane ratio of 1.5:1. Cell adhesion was enabled through the incorporation of 1 mM RGD peptide (GCGTGRGDSPG, Genscript) in all hydrogel groups.

Hydrogel precursor components were mixed and pipetted into the openings of the silicone spacer sheet fitted onto the thiol-functionalized glass piece. Hydrogel precursors were flattened with a glass piece treated with the hydrophobic coating SigmaCote and photopolymerized (365 nm, 5mW/cm^2^) in a UV box (VWR) in the presence of 1 mM lithium acylphosphinate (LAP) photoinitiator for 2 min. The silicone spacer was removed from the glass, and a bottomless 96-well plate was adhered to the functionalized glass piece by exposing the adhesive layer attached to the bottomless plate. Even pressure was applied across the whole plate to ensure a tight seal. PBS was added to wells containing hydrogels and left to swell overnight at 37°C before germicidal UV sterilization and subsequent experimental use. For mechanical characterization, the glass sheet with adhered hydrogels (but no well plate) was added to a large petri dish containing PBS and left to swell overnight at 37°C before nanoindentation. For 3D culture, the same procedure described above was followed with the following differences: the silicone spacer was replaced with the thicker PDMS mold, and a dithiol MMP-degradable peptide (GCKGGPQG↓IWGQGKCG, Genscript) replaced the DTT while maintaining the same thiol:norbornene ratios.

### 2.6 Mechanical characterization

All mechanical characterization was completed via nanoindentation on hydrogels that had been swollen in PBS overnight at 37°C. Indentation tests were performed using Optics11 Life Piuma and Chiaro nanoindenters. A 50 μm diameter borosilicate glass probe attached to a cantilever with a spring constant of 0.5 N/m was used for testing. Indentations were made to a depth of 5 μm using a constant time ramp of 2 sec. The indentation depth was set so that the probe diameter did not exceed 16% of the tip radius to fit Hertzian contact model criteria, and a Poisson’s ratio of 0.5 was assumed. The relaxation response of hydrogels was tested by holding the probe at a fixed depth for 30 sec before lifting the probe. E’ and E’’ were determined using dynamic mechanical analysis (DMA) at frequencies of 0.1-10 Hz and a depth of 5 μm. Surface scans were performed using a matrix of 5 x 5 points with the distance between those points scaled to retain the percentage of the hydrogel surface area that was covered. For hydrogels on a 22 x 22mm coverslip that interval was 1000 μm and for 96-well hydrogels the points were 200 μm apart. During matrix scans, the hydrogel surface was relocated at each point, allowing assessment of the sample surface heterogeneity.

### 2.7 Cell culture

NIH3T3 murine fibroblasts (Sigma-Aldrich) between passage 5-9 were cultured in Gibco Dulbecco’s Modified Eagle Medium (DMEM) supplemented with 10 v/v% fetal bovine serum (FBS) and 1 v/v% antibiotic-antimycotic (1000 U/mL penicillin, 1000 μg/mL streptomycin, and 0.25 μg/mL amphotericin B). Swelled hydrogels were sterilized via germicidal UV irradiation for at least 2 h and incubated in culture medium for at least 30 min prior to cell seeding. Cells were seeded at 1.5 x 10^3^ cells/hydrogel in the 96-well array (6 mm diameter), 2.5 x 10^4^ cells/hydrogel on coverslips (22 x 22 mm), and 1 x 10^6^ cells/mL for 3D culture.

### 2.8 Immunostaining

Cells were fixed on hydrogels using 10% neutral-buffered formalin for 15 min, then permeabilized with 0.1% Triton X-100 in PBS for 10 min. 3 w/v% bovine serum albumin (BSA) in PBS was used to block for at least 1 h at room temperature. Hydrogels were then incubated overnight with primary antibodies against Yes-associated protein (YAP) (mouse monoclonal anti-YAP, 1:400, Sigma-Aldrich) at 4°C. The hydrogels were washed with PBS and incubated with both rhodamine phalloidin to visualize F-actin (1:600, Invitrogen) and secondary antibody (AlexaFluor 488 goat anti-mouse, 1:800) in the dark at room temperature for 2 h. The hydrogels were then rinsed with PBS and stained with DAPI (Invitrogen, 1:10,000) for 1 min before rinsing with PBS.

### 2.9 EdU proliferation assay

Proliferative activity of 2D hydrogel cell cultures was measured using an EdU (5-ethynyl-2’-deoxyuridine) labeling kit (Click-iT™ EdU Cell Proliferation Kit for Imaging, Invitrogen). 5 μM EdU solution was dosed to cells 12 h prior to fixing with 4% paraformaldehyde. Cultures were then permeabilized and stained according to manufacturer’s instructions with 100 μL of stain per well. Plates were then washed and stained with DAPI before proceeding to imaging. EdU staining was done separately from antibody-based staining.

### 2.10 Live/Dead assay

Cell viability of the 3D hydrogel cultures was measured using a fluorescent Live/Dead kit (Invitrogen L3224) following manufacturer instructions. Hydrogels were imaged immediately after the final PBS wash.

### 2.11 Cell fluorescent imaging and analysis

96-well hydrogels were imaged on the Cytation 10 confocal imaging system (Agilent, BioTek) at either 20x or 40x magnification. 2D cell cultures were imaged using beacon-setting and autofocus capabilities. Specifically, for imaging 2D hydrogel cultures, 4 x 4 grids of imaging points (beacons) were applied to each well of interest, and autofocus capabilities were applied onto the DAPI channel at each of these points to find the optimal z-height. 3-slice z-stacks (5 μm slice thickness) were taken at each imaging location with the automatically determined z-height as the middle slice. 3D cell cultures were also imaged using beacon-setting. Specifically, 4 x 4 grids of imaging points were applied to wells of interest. At each of these set x, y beacon locations z-stacks were acquired of 200 μm thickness with 5 μm thick slices. The starting height of these z-stacks was chosen to capture the center height of each of the hydrogels being imaged. Z-stacks from both 2D and 3D imaging experiments were post-processed to generate maximum projection images, and these images were run through deconvolution before running image analysis.

Cell measurements were acquired using a CellProfiler (Broad Institute, Harvard/MIT) pipeline. Cell nuclei were identified using adaptive Otsu thresholding, and then filtered for appropriate circularity to distinguish from any background or debris. When applicable, cell cytoskeletons were then identified as corresponding secondary objects using the same adaptive Otsu thresholding. The integrated intensity of YAP staining was measured across the cell cytoplasm as well as within the nucleus and measurements were normalized to nuclear and cytoplasmic areas. For Live/Dead images, live cells and dead cells were each identified as primary objects using adaptive Otsu thresholding, and the number of live cells identified per image was divided by the total number of cells to obtain viability percentage.

### 2.12 Microplate reader and analysis

For population-level analysis, 96-well hydrogel plates were analyzed on a microplate reader (Tecan Spark Multi-Mode Microplate Reader) immediately following staining. Fluorescent readings were quantified across wells containing hydrogels as well as blank wells with PBS and the gain was optimized for desired runs before measurements were taken. The average fluorescent reading for all blank wells was subtracted from each test well for each respective measurement.

### 2.13 Statistical analysis

For nanoindentation testing, at least three technical hydrogel replicates were tested in which individual point measurements were averaged across each hydrogel and the data were presented as mean ± standard deviation. For cell counting analysis, ordinary one-way ANOVA was first run (not shown) to confirm significant differences between groups with *P* < 0.0001, then ordinary linear regression was run with each slope being significantly different than zero with *P* < 0.0001. For subsequent analysis, statistical tests were chosen based on normality criteria as tested by QQ plots. For data that met normality criteria, ordinary one-way ANOVA was run with Tukey’s post-hoc analysis. For the data that was non-normal, a Kruskal-Wallis test was run with Dunn’s post-hoc comparison. Statistically significant differences are indicated by *, **, or *** corresponding to *P* < 0.05, 0.01, or 0.001 respectively. Details on sample size or additional statistical analysis are included in figure captions.

## 3. Results

### 3.1 Mechanically tunable hyaluronic acid hydrogels are successfully fabricated within 96-well plates

Hyaluronic acid (HA) hydrogels have seen widespread use for cell culture due to their biological relevance and their ability to be highly modified with various functional groups, including a range of crosslinking handle s and bioactive moieties^16^. For example, various modes of HA crosslinking have enabled orthogonal control over mechanical properties^14,17–19^ as well as *in situ* stiffening^20,21^. Further, the independent control of bioactive cues has enabled investigation into the distinct roles of integrin engagement and mechanics on cell behavior^6,22^. Given that HA hydrogels permit a high degree of tunability, there is a need for hydrogel fabrication approaches that allow combinatorial testing of the many independent cell-instructive variables that may be achieved with increased throughput.

In this work, HA was modified with either norbornenes or β-cyclodextrins that enable different modes of crosslinking to achieve a range of stiffnesses and viscoelastic characteristics (**Figure 1A**). The carbon-carbon double bond on norbornene is readily reactive with free thiols through thiol-ene click photochemistry. Thus, with the incorporation of controllable stoichiometric amounts of dithiol crosslinker, a range of covalent crosslinking densities can be achieved to modulate stiffness. Additionally, β-cyclodextrin forms a host-guest complex with adamantane molecules to form physical crosslinks that are reversible under cell-relevant traction forces. The adamantane groups are linked to a cysteine-containing peptide that can be clicked onto the norbornene groups as well. The concentration of adamantane and the ratio of covalent to supramolecular crosslinks in the system are thus very easily tuned (**Figure 1B**).

**Figure 1:**
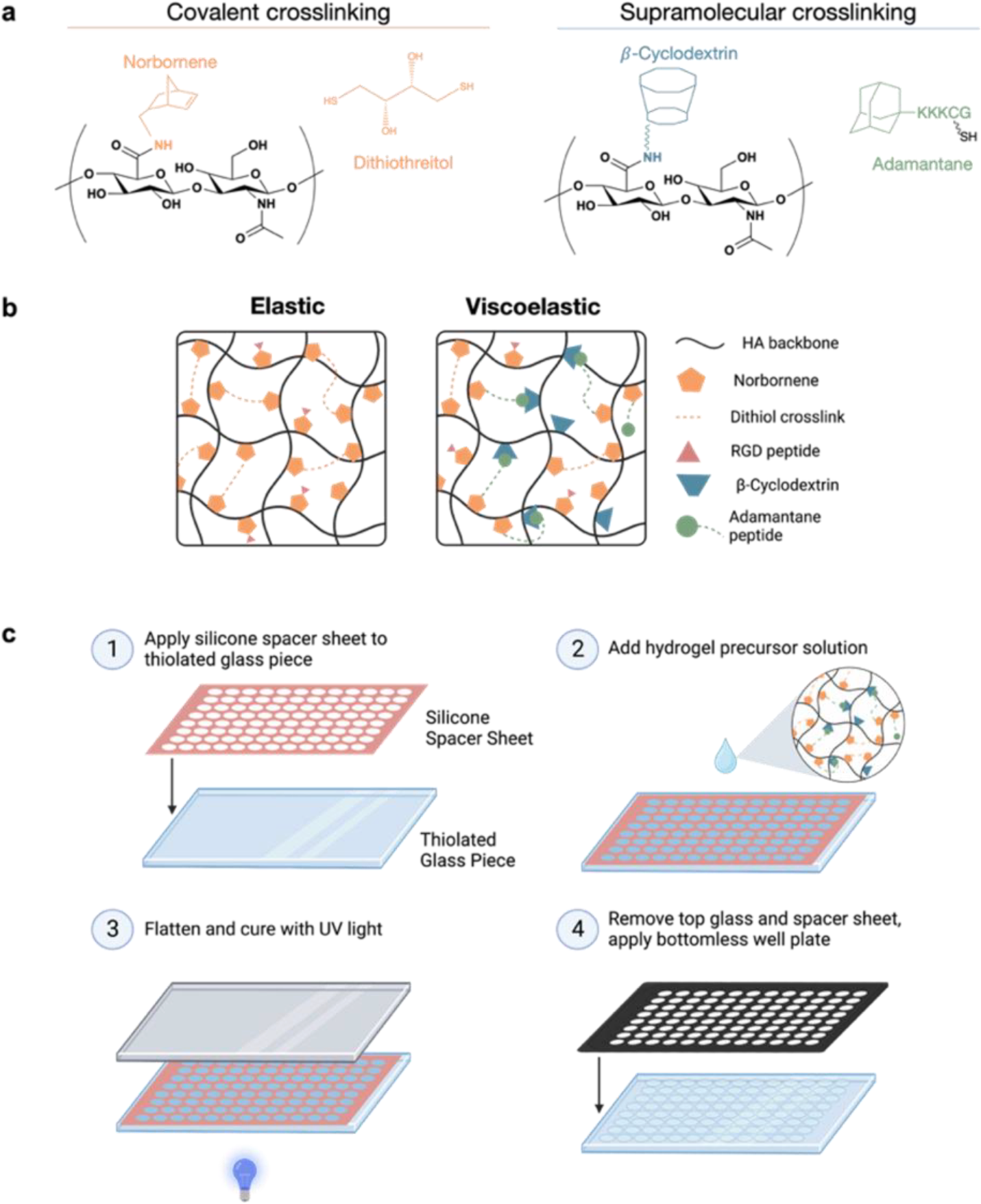
High-throughput fabrication approach to 96-well hyaluronic acid hydrogel array. **a)** Two modes of crosslinking were utilized in this work – covalent and supramolecular – both through the modification of hyaluronic acid (HA). Norbornene-modified HA (NorHA) is covalently crosslinked with dithiothreitol (DTT) through a light-mediated thiol-ene click reaction to form elastic hydrogels. Viscoelastic hydrogels additionally incorporate β-cyclodextrin-adamantane host-guest interactions, where a thiolated adamantane peptide is tethered to the NorHA through the same thiol-ene click reaction. **b)** All components of the elastic or viscoelastic hydrogel networks come together as depicted in the schematic, where hydrogels can also be modified with thiolated RGD peptide to enable cell culture. **c)** 96-well HA hydrogel arrays for 2D cell culture are fabricated through the following steps: 1) functionalization of glass piece with free thiols, and application of silicone spacer sheet onto glass; 2) application of distinct hydrogel precursor solutions into cutouts in the silicone sheet; 3) flattening of hydrogel precursor solutions with SigmaCote-treated glass piece and curing of hydrogels by UV light; and 4) removal of top glass piece and silicone sheet, and application of bottomless 96-well plate.

HA hydrogels have been previously utilized by many labs for two-dimensional (2D) cell culture, typically fabricated on individual coverslips. We modified this existing fabrication method for increased throughput and compatibility with multiwell plate cell culture (**Figure 1C**). Glass pieces sized to the dimension of the 96-well plate were treated with a thiolated silane to introduce free thiols to the substrate surface. Free thiols could then react with the norbornenes on the HA ranging up to 4 wt% can span elastic moduli of two orders of magnitude, while keeping the fraction of norbornenes crosslinked the same (**Figure 2A**). This is to be expected, given that a higher weight percent of polymer has a greater concentration of norbornenes as well as an increase in total polymer content. Additionally, hydrogels of the same polymer weight percent were fabricated with different crosslinking densities to achieve various stiffnesses. Increasing crosslink density in hydrogels of a lower weight percent had less of an influence on stiffness than increasing the crosslink density in hydrogels of a higher weight percent. Through the variation of both the polymer weight percentage and the percent of norbornene crosslinks, the stiffness of the resultant hydrogels can be precisely tuned.

**Figure 2:**
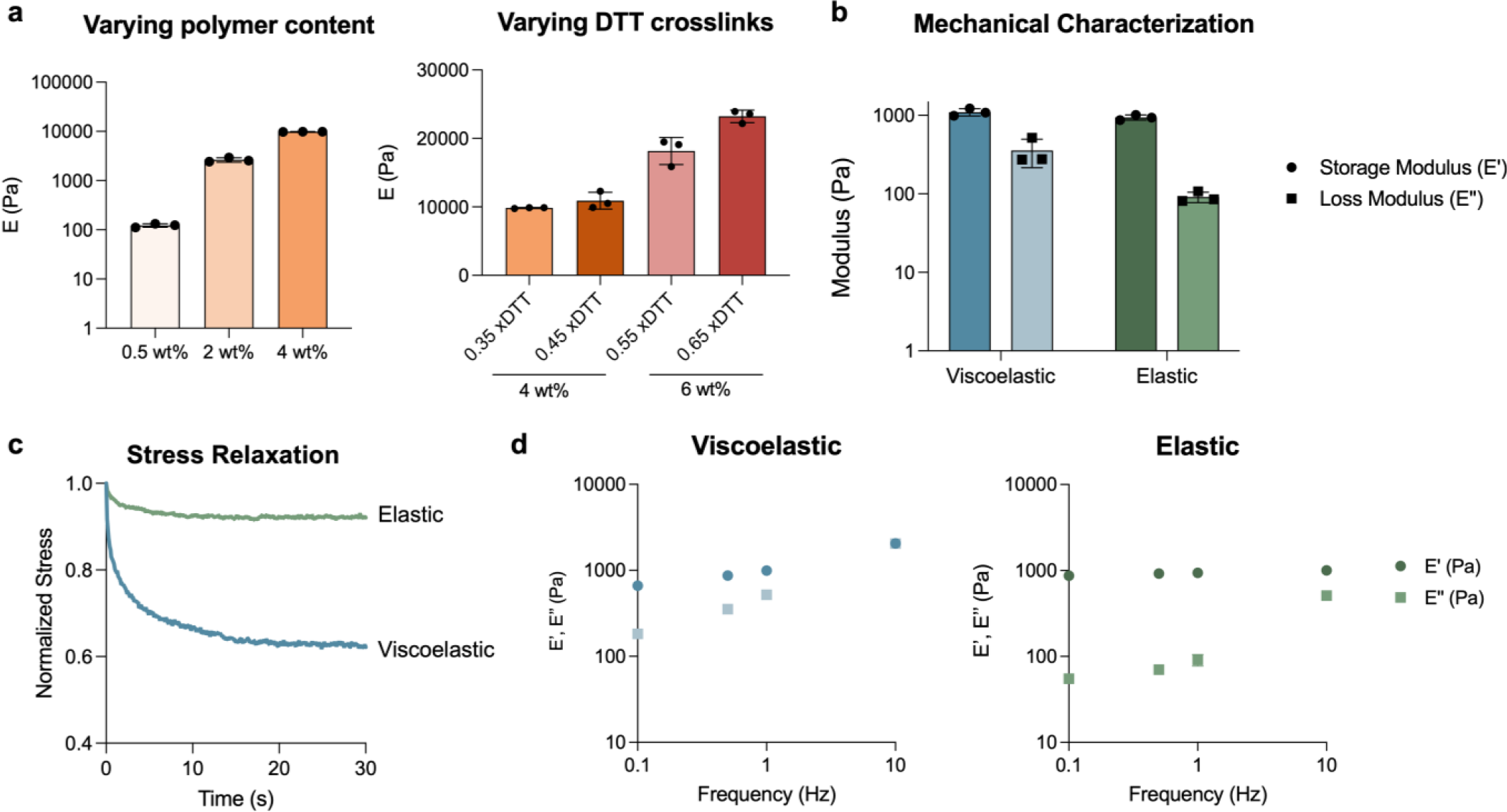
*In situ* mechanical characterization of 96-well hydrogel array. **a)** NorHA hydrogels fabricated within a 96-well plate were swollen overnight in PBS and mechanically characterized *in situ* using nanoindentation. NorHA hydrogels of varying polymer content spanned a tissue-relevant stiffness range of three orders of magnitude. Hydrogels in each group had a thiol:norbornene ratio of 0.35. Additionally, changing the thiol:norbornene ratio while maintaining the polymer weight percent produced a more gradual change in stiffness. **b)** Viscoelastic and elastic hydrogels of matching storage moduli were fabricated within the same well plate. The loss modulus of the viscoelastic formulation was confirmed to be within one order of magnitude of the storage modulus. This was determined using *in situ* DMA testing at 1 Hz. **c)** Hydrogels of stiffness-matched formulations were tested for stress relaxation behavior by nanoindentation. Viscoelastic hydrogels demonstrated a greater extent of relaxation than elastic hydrogels. **d)** Stiffness-matched elastic and viscoelastic hydrogels were also tested for storage and loss moduli at varying frequencies using DMA testing. Viscoelastic hydrogels demonstrated increased moduli with increasing frequency, whereas elastic hydrogels did not show frequency-dependent behavior at lower frequencies. *n* = 3 hydrogels per group.

The viscoelastic characteristics of hydrogels in the 96-well plate can be tuned through the introduction of supramolecular crosslinking. Dynamic mechanical analysis (DMA) was performed using the same nanoindentation approach on swollen hydrogels to measure the complex modulus of hydrogels *in situ*. The number of physical and covalent crosslinks was tuned to fabricate elastic or viscoelastic hydrogels of equal storage modulus, but with loss moduli over an order of magnitude different (**Figure 2B**). Additionally, viscoelastic characteristics were investigated through stress relaxation tests and frequency sweeps. As seen in **Figure 2C**, hydrogels fabricated with physical crosslinks showed a much greater extent of stress relaxation after 30 seconds than hydrogels fabricated with covalent crosslinks only. Additionally, DMA showed that viscoelastic hydrogels had increasing moduli with increasing frequency, whereas covalently crosslinked elastic hydrogels showed little frequency dependence at lower frequencies (**Figure 2D**). Taken together, we can conclude that the incorporation of physical crosslinks into these hydrogels imparts viscoelastic and stress relaxing character.

Once we confirmed that we could fabricate hydrogels with varying stiffness and viscoelasticity within the same 96-well array, we sought to quantify changes in fabrication throughput compared to our conventional approach of fabricating individual hydrogels on coverslips. Each fabrication step was timed independently and across at least two different researchers. The fabrication time required for each step was divided by the total number of hydrogels fabricated and this was used to estimate the total fabrication time required to generate from 1 to 96 individual hydrogels either in a 96-well plate or on coverslips. The total fabrication time was reduced by 2-fold for 3 hydrogels and over 8-fold for a full 96-well plate (**Figure 3A**). The reduction of hydrogel volume from 55 μL to 18 μL in the 96-well plate format resulted in a uniform 3-fold reduction in the amount of polymer material needed to fabricate the same number of hydrogels on coverslips (**Figure 3B**). Overall, this fabrication approach enables significant time and material savings for small-scale experiments while also enabling higher throughput on large-scale experiments.

**Figure 3:**
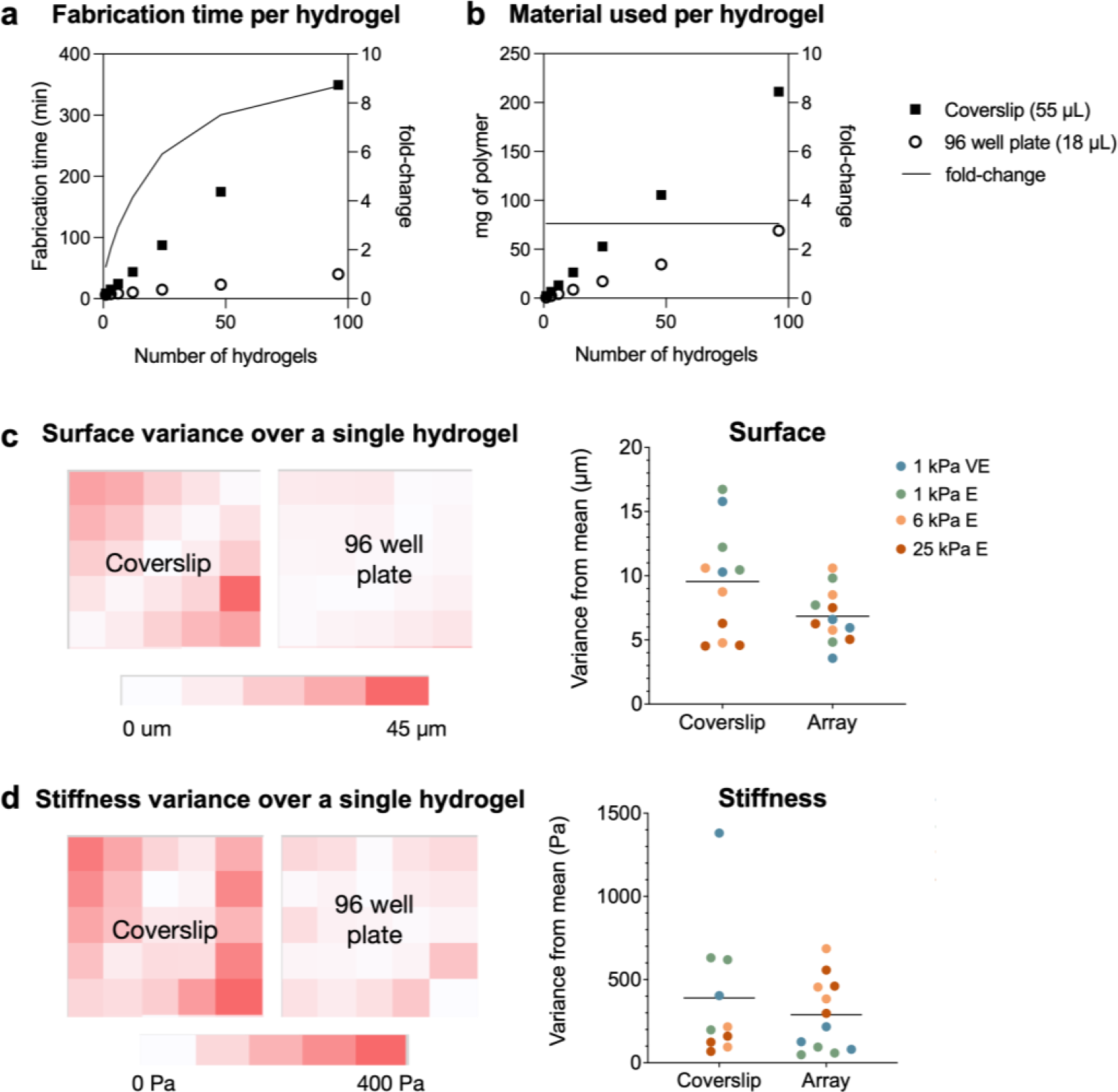
Comparison of hydrogel characteristics fabricated within 96-well plates or on coverslips. **a)** The time required for each step in hydrogel fabrication was measured across five experiments with two independent researchers. The time for each step was averaged across runs, and steps that were dependent on the number of replicates were reduced to the amount of time per replicate. These time values were then used for extrapolation to the total fabrication time required per replicate in either fabrication format. The 96-well plate fabrication approach enables up to an 8-fold reduction in fabrication time per hydrogel. **b)** The amount of material needed for a 4 wt% hydrogel was calculated to be 3-fold less on a per hydrogel basis in the 96-well array versus coverslip format. **c)** *In situ* nanoindentation was used to evaluate surface heterogeneity across three hydrogels of varying mechanical characteristics. The step size between test points on the coverslip hydrogels and 96-well hydrogels was 1000 μm and 400 μm respectively, to evaluate near the same percentage of the total hydrogel surface area. The absolute variation from the mean surface height was averaged for each hydrogel and plotted. Across all mechanical groups tested, there was comparable surface flatness. **d)** The same analysis was then performed to assess stiffness heterogeneity using *in situ* nanoindentation. Again, there was no change in mechanical heterogeneity with the adaptation of the 96-well plate fabrication approach.

The characteristics of the resultant hydrogels were also investigated to ensure that the 96-well array fabrication approach maintained expected hydrogel characteristics. Surface flatness was characterized by nanoindentation using swollen hydrogels of identical formulations fabricated either within a 96-well plate or on a 22 x 22 mm coverslip. An equal area was investigated on each type of hydrogel and the variance from the average height was visualized as a heat map or plotted for each hydrogel (**Figure 3C**). Similarly, stiffness uniformity was also investigated through nanoindentation and plotted in the same manner (**Figure 3D**). For both surface flatness and stiffness, the 96-well plate fabrication approach showed statistically similar variance from the mean across the same area. These results indicate that hydrogels generated within 96-well plates do not lose appropriate uniformity characteristics despite the increased fabrication throughput.

### 3.2 96-well plate hydrogel platform is amenable to high-throughput cellular analysis at both the population and single cell-level

Following the validation of successful 96-well hydrogel platform fabrication, we sought to investigate which high-throughput approaches would be suitable for taking population-level and individual cell-level measurements. As 96-well plates are compatible with microplate reader devices, we measured changes in cellular abundance using this approach. We fabricated a 96-well plate containing hydrogels of identical mechanics (6 kPa, elastic) and seeded NIH3T3 fibroblasts at a backbone to covalently bond hydrogels to the glass through thiol-ene click chemistry. Following glass treatment, a silicone sheet was applied onto the glass, and hydrogel precursor solution was added into the inlet regions of the mold. A hydrophobic glass piece was then applied on top to achieve a uniform culture surface, and UV light was applied to trigger hydrogel crosslinking. The top glass piece and silicone mold were then removed, and the glass substrate was used either for *in situ* mechanical indentation testing, or a bottomless 96-well plate was applied for subsequent cell culture. The suitability of the adhesive-bound well plate for cell culture was assessed by monitoring liquid leaking between wells, and no spillover was observed in a 24 hour period (**Figure S9, Supplementary Movie 1**). Thus, the application of the 96-well plate following hydrogel formation was determined to be a suitable approach to enabling both *in situ* mechanical characterization as well as isolated culture in individual wells.

The modular nature of this fabrication approach allows for easy adaptability of various hydrogel precursor formulations. As an illustrative example, we fabricated a hydrogel array where HA polymer content and crosslink density were varied to achieve a range of mechanical properties. Following fabrication of the array, hydrogels were swollen overnight in PBS, and the resultant mechanical properties were evaluated using nanoindentation. The *in situ* characterization of hydrogels in the array is advantageous, as hydrogel mechanics are known to vary with specimen geometry^23^. HA hydrogels of 0.5 wt% range of densities. The cultures were fixed and stained using a standard approach often employed for hydrogel-based cell culture experiments, and we compared microplate reader measurements to quantification determined via automatic imaging or manual imaging. All three methods were able to detect a strong correlation in the increase in nuclear (DAPI) signal with the increase in number of cells seeded (**Figure 4A**). The microplate reader showed the best agreement with this positive correlation, indicating that this method is useful for detecting trends in cellular behavior at a population-level.

**Figure 4:**
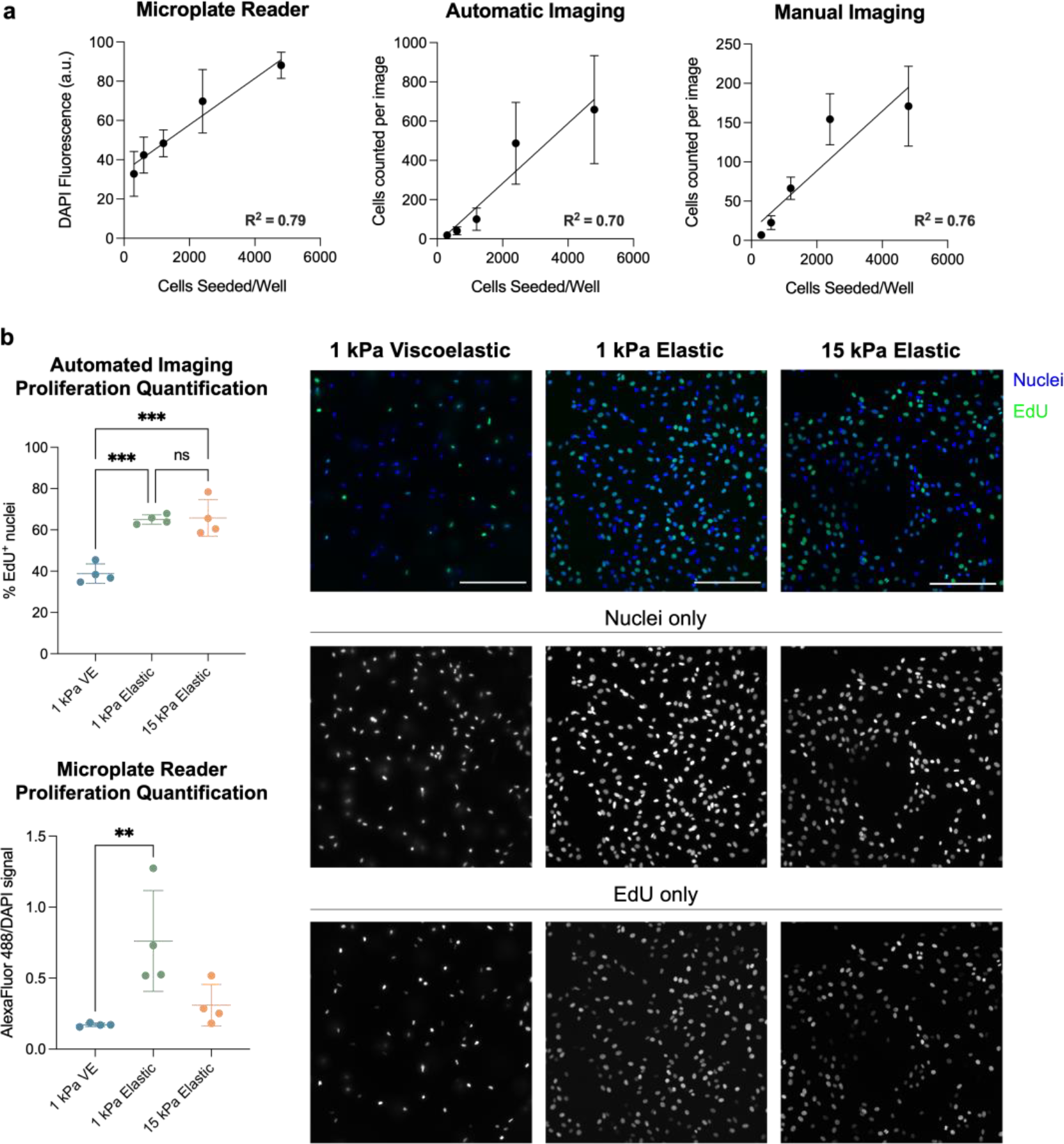
Evaluation of high-throughput methods for cellular phenotype evaluation at the population-level. **a)** NIH3T3 fibroblasts were seeded on 6 kPa elastic hydrogels within 96-well plates and cultured for 3 days. Following fluorescent staining of cell nuclei and F-actin, cell counts were performed by either quantifying DAPI fluorescence on a microplate reader, manual imaging, or imaging using an automated pipeline. Simple linear regression was run for each dataset and signal from the microplate reader showed the best correlation with the number of cells seeded per well. Representative images were taken using manual imaging. Data are plotted as the mean +/-SD for *n* = 5 hydrogel replicates per group. **b)** NIH3T3 fibroblasts were seeded on 1 kPa elastic or viscoelastic and 15 kPa elastic hydrogels and treated with EdU labeling solution 12 hours before fixing (*n* = 4 hydrogels). Image quantification was able to detect significant differences in EdU positivity between the viscoelastic and elastic groups, whereas the microplate reader quantification was less sensitive to these differences. Images were taken using an automated imaging protocol. Image quantification data was analyzed using an ordinary one-way ANOVA. Microplate reader quantification was non-normal and thus analyzed using a Kruskal-Wallis test. *\*\*: P* < 0.01, ***: *P* < 0.001. Scale bars = 200 μm.

We then sought to investigate the sensitivity of plate reader measurements to distinguish cellular behaviors within the 96-well plate hydrogel platform. We seeded NIH3T3 fibroblasts on hydrogels of varying stiffnesses and investigated how proliferative activity changes between hydrogel mechanical environments using EdU staining. As would be expected^24–26^, cells on elastic substrates show higher proliferative activity than those on viscoelastic substrates (**Figure 4B**). These differences were easily identified through high-content automatic imaging and subsequent analysis. While the same trends remained visible with the microplate reader, this modality showed less of an ability to distinguish incremental changes in cellular phenotype at this scale. Thus, this data suggests that although the 96-well plate hydrogel platform is amenable to high-throughput measurements acquired by both the microplate reader and automated imager, the microplate reader is not well suited for detecting modest changes in cellular phenotype in this platform.

Next, we investigated whether the 96-well hydrogel array was suitable for the evaluation of cellular behavior on a single cell level. Cell-level measurements are of particular interest when investigating cell response to substrate mechanics, given that many phenotypic changes are both sub-cellular and heterogeneous, and therefore not distinguishable from population-level analyses. We chose to investigate cell spreading, protein nuclear localization, and F-actin organization to demonstrate this platform’s ability to support quantification of single-cell metrics. Specifically, we sought to optimize a protocol for successful high-magnification imaging across variable hydrogel nodes that maintained sufficient image quality to distinguish sub-cellular changes in protein organization. NIH3T3 fibroblasts were seeded on hydrogels of varying mechanics and cultured for 3 days before fixing and staining. Images were acquired using automatic focusing and acquisition of pre-programmed locations in each well (**Figure 5A**). These images were analyzed for a variety of cellular and sub-cellular metrics. Although cell spread area did not vary significantly between mechanical groups, the nuclear localization of the mechanoresponsive transcriptional coactivator Yes-associated protein (YAP) significantly increased in cells cultured on stiffer hydrogels (**Figure 5B**). Additionally, there were significant changes in the angular second moment, a metric quantifying uniformity, of the F-actin cytoskeleton in the elastic groups compared to the viscoelastic group. This decrease in angular second moment is likely indicative of an increase in F-actin organization into stress fibers that form when fibroblasts are mechanically activated^27^. Altogether, these data demonstrate that this 96-well plate hydrogel platform enables the high-throughput acquisition of rich cell-level data describing changes in cellular phenotype due to mechanical cues.

**Figure 5:**
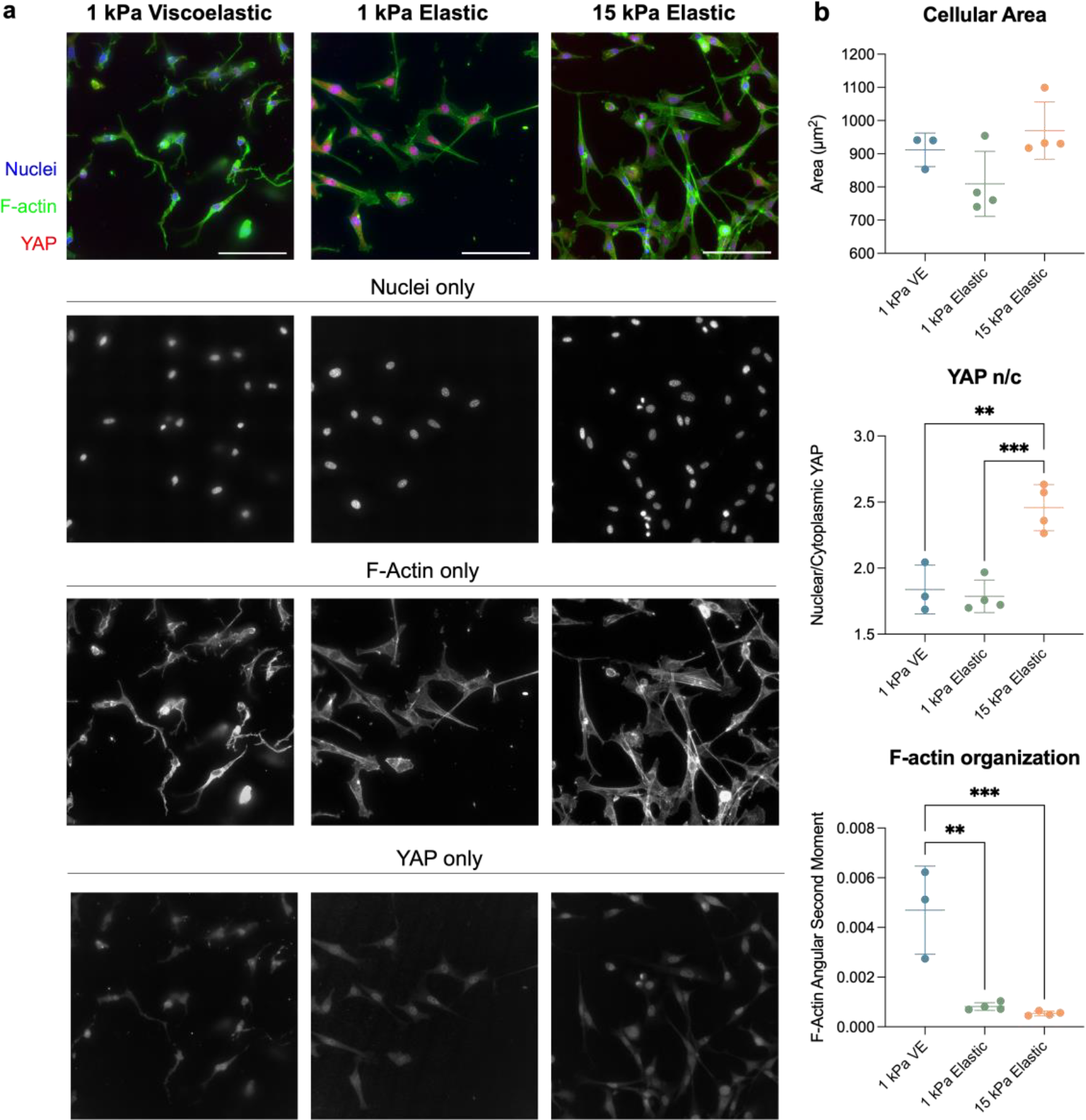
Automated image acquisition and analysis for evaluating of cell response to 96-well hydrogel array mechanical cues. **a)** Representative images of NIH3T3 cells cultured on 1 kPa viscoelastic or elastic hydrogels and 15 kPa elastic hydrogels. **b)** Cell spread area was similar for all groups, but YAP nuclear localization was significantly increased in the 15 kPa group. Additionally, an increase in F-actin organization, denoted by lower angular second moment, was observed in the elastic groups. *n* = 4 hydrogels per group. All cellular metrics were analyzed using an ordinary one-way ANOVA. *\*\*: P* < 0.01, ***: *P* < 0.001. Scale bars = 100 μm.

### 3.3 96-well plate hydrogel platform is compatible with 3D cell culture

Finally, we sought to explore the capabilities of our 96-well plate hydrogel platform for 3D cell culture applications. The aforementioned hydrogel fabrication approach was easily adaptable for 3D cell culture (**Figure 6A**). Specifically, a 1 mm thick PDMS mold of similar geometry to the silicone spacer used in 2D culture was utilized to increase the height of the hydrogels. This PDMS mold was then applied to the thiolated glass piece and cells suspended in hydrogel precursor solutions were pipetted into the appropriate geometries. No flattening step was required for 3D hydrogel formation, and when the PDMS mold was removed the cell-laden hydrogels remained tethered to the glass substrate. A bottomless 96-well plate was then adhered, and media was added onto the hydrogels for subsequent cell culture. This approach for 3D cell culture within 96-well plates requires no additional time or equipment, making this an equally accessible approach for answering many biological questions with reasonable throughput and biological relevance.

**Figure 6:**
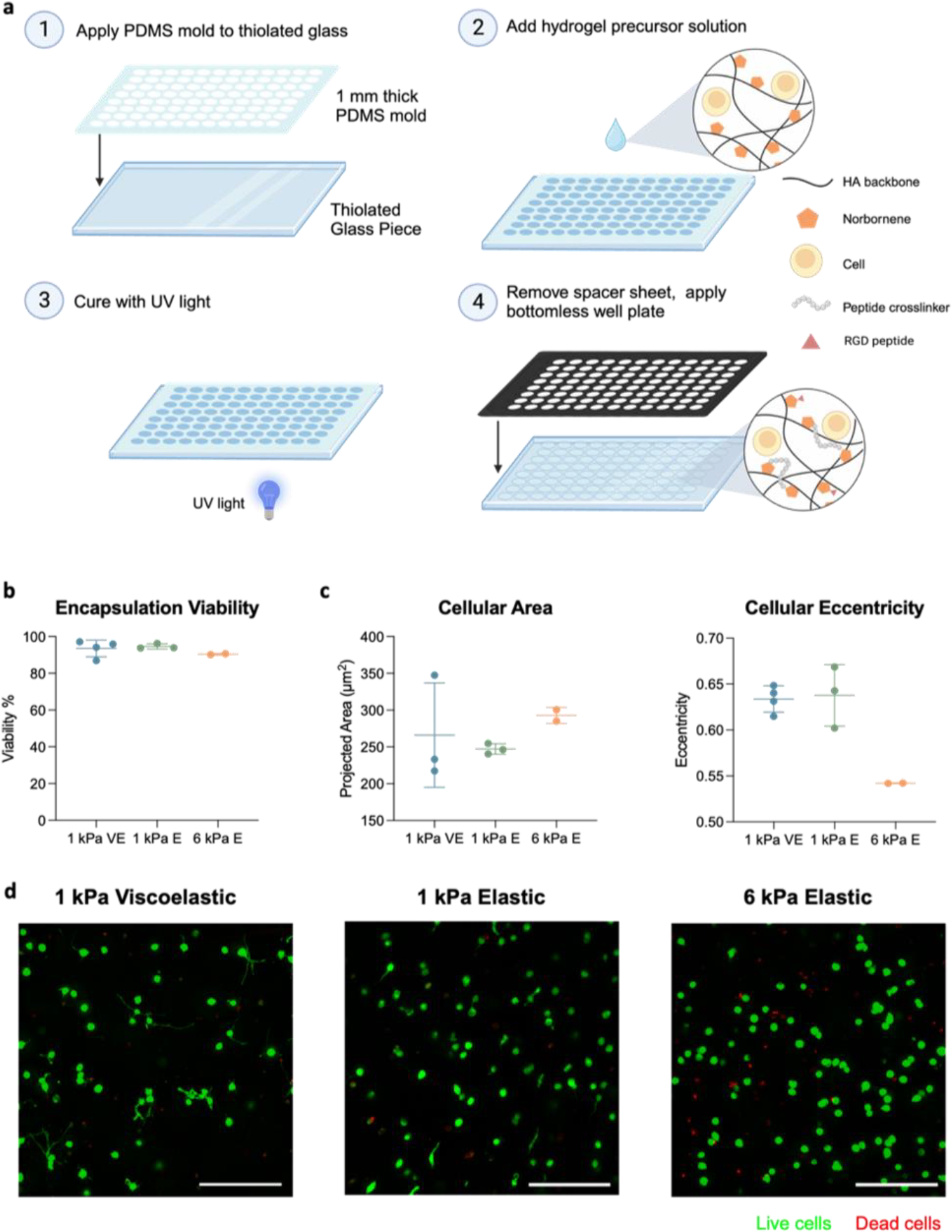
3D cell encapsulation within the 96-well hydrogel array. **a)** 96-well hydrogel arrays for 3D cell culture were fabricated similarly to those used for 2D culture with the following steps: 1) functionalization of glass piece with free thiols, and application of PDMS mold onto glass; 2) application of cell-laden hydrogel precursor solutions into cutouts in the PDMS; 3) curing of hydrogels by UV light; and 4) removal of PDMS mold, and application of bottomless 96-well plate. **b)** NIH3T3 cells encapsulated in 1 kPa viscoelastic or elastic hydrogels, or 6 kPa elastic hydrogels all showed high viability (> 90%) after 2 days of culture. **c)** Live cell morphology was analyzed using CellProfiler, and cell projected area trended higher with increased stiffness. Additionally, cells had fewer protrusions and tended to be more circular in the 6 kPa hydrogels. **d)** Representative images of live (*green*) and dead (*red*) cells acquired using automated imaging are shown. Scale bars = 200 μm.

We then investigated whether this platform would support high cell viability and remain amenable to cell-level phenotypic measurements. We fabricated both viscoelastic and elastic 3D hydrogels with the same norbornene-modified HA, replacing all DTT covalent crosslinker with MMP-degradable peptide crosslinkers. We encapsulated NIH3T3 fibroblasts in these hydrogels within the 96-well plate and cultured them for 2 days before staining for viability using a Live/Dead assay and imaging on a high-content confocal microscope. All groups showed high viability (> 90%, **Figure 6B**). Additionally, cells within each experimental group showed distinct morphological behavior. The greatest number of cellular protrusions was seen in the viscoelastic group, whereas the cells in the 6 kPa elastic group were the most rounded with higher projected cell areas (**Figure 6C**). Taken together, these results demonstrate the utility of this platform for the acquisition of robust cellular-level data in 3D culture.

## 4. Discussion

As the field of hydrogel development continues to progress, there is a need to enhance the accessibility of this growing set of tools. Hydrogel cell culture models are often limited by the number of variables that can be investigated simultaneously due to the time and material requirements for these experiments. Additionally, commercially available hydrogel plates, such as CytoSoft^®^, indicate a growing interest in the adaptation of hydrogel culture systems outside the biomaterials field. Yet, these plates come at a large expense and suffer from a lack of modularity or mechanical complexity. Thus, there is an opportunity for accessible hydrogel fabrication approaches that work to enable experimentation for those both new to and seasoned in the hydrogel field.

Although we utilized thiol-ene hyaluronic acid hydrogels in this work, the fabrication approach described here is adaptable to many different material systems. Specifically, any polymer backbone that will support these polymer modifications could easily be substituted, such as poly(ethylene glycol) (PEG), alginate, or gelatin^28–30^. Further, the specific alkene modification employed here (norbornene-modified HA) may be substituted for other widely used alkene modifications such as vinyl sulfone, maleimide, and (meth)acrylate groups^31–37^. Thus, this approach may be leveraged to increase the throughput of many recently developed hydrogel technologies, such as the ability to present and remove unique biological cues in each well^38–40^, to induce differential amounts of photodegradation between wells^41,42^, or to induce stiffening over time through subsequent curing^20^. Additionally, this approach is not limited to thiol-ene chemistries but can be extended to any silane-modified surface functionalization that enables the simultaneous bonding and glass surface-grafting of the polymer network of interest. Ultimately, this approach is highly versatile, and easily adapted to a wide range of hydrogel chemistries, whether commercially available or synthesized in house.

Additionally, multiwell plate geometries beyond 96-wells are amenable to this fabrication approach. Bottomless well plates are available for plates spanning 6-wells to 384-wells. Thus, for experiments that require higher cell numbers at the endpoint of the culture period— such as Western blots, qPCR, or RNA sequencing— this approach may be adapted to 12 or 24 well platforms. For applications with higher throughput needs, such as drug screening, hydrogels within 384-well plates can be utilized, although this level of throughput may be challenging to use without automation. Yet many microplate readers and high-content imaging systems, including those used in this study, are easily adapted to 384-well plate geometries, making this an attractive approach for these types of studies as has been shown by others^12^.

Finally, the adaptation of this approach for 3D cell culture is a powerful tool in moving toward more physiologic culture conditions^43^. Yet, the adherence of 3D hydrogels in this platform offers both advantages and limitations. The advantages of this approach include the simplicity in fabrication, the ease of media transfers, and the ease of image acquisition. However, the fixation of the hydrogel surface to the glass may restrict swelling and/or nutrient diffusion. Although we saw no limitations in cell viability in the culture conditions tested, this fabrication approach may hinder successful culture for sensitive cell types or longer-term cultures.

## 5. Conclusion

In this work we demonstrated the successful fabrication of 96-well hyaluronic acid hydrogel arrays to enable increased experimental throughput in both 2D and 3D cell culture studies. We utilized thiol-ene chemistry to fabricate hydrogels of stiffnesses across three orders of magnitude and integrated host-guest supramolecular interactions in hydrogel networks to impart viscoelastic characteristics. These hydrogels demonstrated comparable surface properties and mechanical homogeneity to the current standard coverslip fabrication method while enabling a 3-fold reduction in the material necessary per hydrogel replicate and up to an 8-fold reduction in time required for fabrication. These hydrogel platforms also successfully supported both population-level and cell-level phenotypic measurements through high-content imaging and microplate readings. Microplate reader analysis was shown to be useful for identifying trends in phenotypic behavior across groups. High-content imaging was also shown to be a powerful tool for generating rich single cell-level data from hydrogel cultures. Imaging metrics were able to detect differences in YAP nuclear localization as well as differences in F-actin organization. Overall, this platform represents an accessible and versatile approach for fabricating hydrogels with higher throughput for cell culture applications.

## Supporting Information

^1^H NMR spectra for the hydrogel components and design schematics for fabricating the 96-well hydrogel array can be found in the Supporting Information.

## Acknowledgments

The authors would like the thank Dr. Jenna Sumey for providing the norbornene-modified hyaluronic acid utilized in this work, Dr. Christopher Highley for providing training, materials, and discussion for the successful completion of 3D printing and PDMS mold casting, Kelly Bukovic for providing training and assistance with ^1^H NMR and for helpful discussions, and Olivia Morren for helpful feedback on writing. We would also like to acknowledge Dwight Dart and the MAE

Rapid Prototyping Lab at UVA for their assistance with 3D printing, as well as Trevor Kemp and UVA School of Architecture’s FabLab for their training and assistance with laser cutting. This work was supported by the NIH (R35GM138187) and NSF (CAREER DMR/BMAT 2046592, GRFP to M.L.S. and L.R.A.). The content is solely the responsibility of the authors and does not necessarily represent the official views of the National Institutes of Health.

## Notes

### Competing Interest Statement

The authors have declared no competing interest.

## References

1. Engler, A. et al. Substrate compliance versus ligand density in cell on gel responses. Biophys J 86, 617–628 (2004).

2. Huang, X. et al. Matrix Stiffness–Induced Myofibroblast Differentiation Is Mediated by Intrinsic Mechanotransduction. Am J Respir Cell Mol Biol 47, 340–348 (2012).

3. Balestrini, J. L., Chaudhry, S., Sarrazy, V., Koehler, A. & Hinz, B. The mechanical memory of lung myofibroblasts. Integr Biol (Camb) 4, 410–421 (2012).

4. Chaudhuri, O. et al. Hydrogels with tunable stress relaxation regulate stem cell fate and activity. Nature Mater 15, 326–334 (2016).

5. Chaudhuri, O. et al. Substrate stress relaxation regulates cell spreading. Nat Commun 6, 6365 (2015).

6. Hui, E., Moretti, L., Barker, T. H. & Caliari, S. R. The Combined Influence of Viscoelastic and Adhesive Cues on Fibroblast Spreading and Focal Adhesion Organization. Cel. Mol. Bioeng. 14, 427–440 (2021).

7. Wipff, P.-J., Rifkin, D. B., Meister, J.-J. & Hinz, B. Myofibroblast contraction activates latent TGF-β1 from the extracellular matrix. Journal of Cell Biology 179, 1311–1323 (2007).

8. Arora, P. D., Narani, N. & McCulloch, C. A. G. The Compliance of Collagen Gels Regulates Transforming Growth Factor-β Induction of α-Smooth Muscle Actin in Fibroblasts. The American Journal of Pathology 154, 871–882 (1999).

9. Grubb, M. L. & Caliari, S. R. Fabrication approaches for high-throughput and biomimetic disease modeling. Acta Biomater 132, 52–82 (2021).

10. Lei, R. et al. Multiwell Combinatorial Hydrogel Array for High-Throughput Analysis of Cell–ECM Interactions. ACS Biomater. Sci. Eng. 7, 2453–2465 (2021).

11. Brooks, E. A., Jansen, L. E., Gencoglu, M. F., Yurkevicz, A. M. & Peyton, S. R. Complementary, Semiautomated Methods for Creating Multidimensional PEG-Based Biomaterials. ACS Biomater. Sci. Eng. 4, 707–718 (2018).

12. Mih, J. D. et al. A Multiwell Platform for Studying Stiffness-Dependent Cell Biology. PLoS One 6, e19929 (2011).

13. Gramlich, W. M., Kim, I. L. & Burdick, J. A. Synthesis and orthogonal photopatterning of hyaluronic acid hydrogels with thiol-norbornene chemistry. Biomaterials 34, 9803–9811 (2013).

14. Hui, E., Gimeno, K. I., Guan, G. & Caliari, S. R. Spatiotemporal Control of Viscoelasticity in Phototunable Hyaluronic Acid Hydrogels. Biomacromolecules 20, 4126–4134 (2019).

15. Rodell, C. B., Kaminski, A. L. & Burdick, J. A. Rational Design of Network Properties in Guest–Host Assembled and Shear-Thinning Hyaluronic Acid Hydrogels. Biomacromolecules 14, 4125–4134 (2013).

16. Highley, C. B., Prestwich, G. D. & Burdick, J. A. Recent advances in hyaluronic acid hydrogels for biomedical applications. Current Opinion in Biotechnology 40, 35–40 (2016).

17. Lou, J., Stowers, R., Nam, S., Xia, Y. & Chaudhuri, O. Stress relaxing hyaluronic acid-collagen hydrogels promote cell spreading, fiber remodeling, and focal adhesion formation in 3D cell culture. Biomaterials 154, 213–222 (2018).

18. Wang, H. et al. Covalently Adaptable Elastin-Like Protein– Hyaluronic Acid (ELP–HA) Hybrid Hydrogels with Secondary Thermoresponsive Crosslinking for Injectable Stem Cell Delivery. Advanced Functional Materials 27, 1605609 (2017).

19. Nelson, B. R. et al. Photoinduced Dithiolane Crosslinking for Multiresponsive Dynamic Hydrogels. Advanced Materials n/a, 2211209.

20. Guvendiren, M. & Burdick, J. A. Stiffening hydrogels to probe short- and long-term cellular responses to dynamic mechanics. Nat Commun 3, 792 (2012).

21. Sumey, J. L., Johnston, P. C., Harrell, A. M. & Caliari, S. R. Hydrogel mechanics regulate fibroblast DNA methylation and chromatin condensation. Biomater. Sci. 11, 2886–2897 (2023).

22. Cosgrove, B. D. et al. N-cadherin adhesive interactions modulate matrix mechanosensing and fate commitment of mesenchymal stem cells. Nature Mater 15, 1297–1306 (2016).

23. Oyen, M. L. Mechanical characterisation of hydrogel materials. International Materials Reviews 59, 44–59 (2014).

24. Yeh, Y.-T. et al. Matrix Stiffness Regulates Endothelial Cell Proliferation through Septin 9. PLOS ONE 7, e46889 (2012).

25. Umesh, V., Rape, A. D., Ulrich, T. A. & Kumar, S. Microenvironmental Stiffness Enhances Glioma Cell Proliferation by Stimulating Epidermal Growth Factor Receptor Signaling. PLOS ONE 9, e101771 (2014).

26. Sun, M. et al. Effects of Matrix Stiffness on the Morphology, Adhesion, Proliferation and Osteogenic Differentiation of Mesenchymal Stem Cells. Int J Med Sci 15, 257–268 (2018).

27. Nelson, A. R., Christiansen, S. L., Naegle, K. M. & Saucerman, J. J. Logic-based mechanistic machine learning on highcontent images reveals how drugs differentially regulate cardiac fibroblasts. 2023.03.01.530599 Preprint at 10.1101/2023.03.01.530599 (2023).

28. Lin, C.-C., Ki, C. S. & Shih, H. Thiol–norbornene photoclick hydrogels for tissue engineering applications. Journal of Applied Polymer Science 132, (2015).

29. Fairbanks, B. D. et al. A Versatile Synthetic Extracellular Matrix Mimic via Thiol-Norbornene Photopolymerization. Adv Mater 21, 5005–5010 (2009).

30. Münoz, Z., Shih, H. & Lin, C.-C. Gelatin hydrogels formed by orthogonal thiol–norbornene photochemistry for cell encapsulation. Biomater. Sci. 2, 1063–1072 (2014).

31. Azagarsamy, M. A. & Anseth, K. S. Bioorthogonal Click Chemistry: An Indispensable Tool to Create Multifaceted Cell Culture Scaffolds. ACS Macro Lett. 2, 5–9 (2013).

32. Hoyle, C. E. & Bowman, C. N. Thiol–Ene Click Chemistry. Angewandte Chemie International Edition 49, 1540–1573 (2010).

33. Nair, D. P. et al. The Thiol-Michael Addition Click Reaction: A Powerful and Widely Used Tool in Materials Chemistry. Chem. Mater. 26, 724–744 (2014).

34. Kharkar, P. M., Rehmann, M. S., Skeens, K. M., Maverakis, E. & Kloxin, A. M. Thiol–ene Click Hydrogels for Therapeutic Delivery. ACS Biomater. Sci. Eng. 2, 165–179 (2016).

35. Hui, E., Sumey, J. L. & Caliari, S. R. Click-functionalized hydrogel design for mechanobiology investigations. Mol. Syst. Des. Eng. 6, 670–707 (2021).

36. Li, X. & Xiong, Y. Application of “Click” Chemistry in Biomedical Hydrogels. ACS Omega 7, 36918–36928 (2022).

37. A. Sawicki, L. & M. Kloxin A. Design of thiol–ene photoclick hydrogels using facile techniques for cell culture applications. Biomaterials Science 2, 1612–1626 (2014).

38. DeForest, C. A. & Anseth, K. S. Photoreversible Patterning of Biomolecules within Click-Based Hydrogels. Angewandte Chemie 124, 1852–1855 (2012).

39. Gould, S. T., Darling, N. J. & Anseth, K. S. Small peptide functionalized thiol–ene hydrogels as culture substrates for understanding valvular interstitial cell activation and de novo tissue deposition. Acta Biomaterialia 8, 3201–3209 (2012).

40. Grim, J. C. et al. A Reversible and Repeatable Thiol–Ene Bioconjugation for Dynamic Patterning of Signaling Proteins in Hydrogels. ACS Cent. Sci. 4, 909–916 (2018).

41. Ki, C. S., Shih, H. & Lin, C.-C. Facile preparation of photodegradable hydrogels by photopolymerization. Polymer 54, 2115–2122 (2013).

42. Wu, L., Di Cio, S., Azevedo, H. S. & Gautrot, J. E. Photoconfigurable, Cell-Remodelable Disulfide Cross-linked Hyaluronic Acid Hydrogels. Biomacromolecules 21, 4663–4672 (2020).

43. Caliari, S. R. & Burdick, J. A. A practical guide to hydrogels for cell culture. Nat Methods 13, 405–414 (2016).

