## Supplementary Information for "Modular multiwell viscoelastic hydrogel platform for two- and three-dimensional cell culture applications"

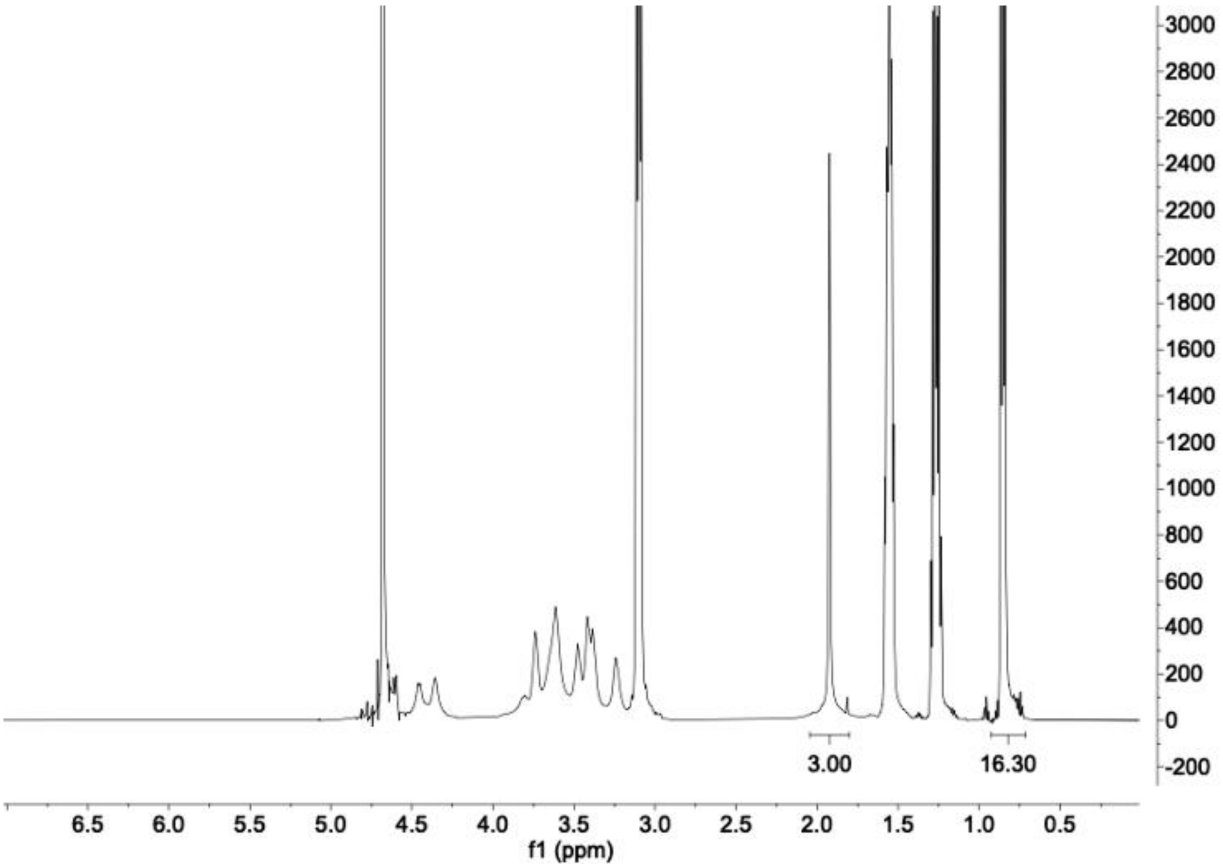


**Supplementary Figure 1: $^1$H NMR spectrum of tetrabutyl ammonium (TBA) salt of hyaluronic acid (HA-TBA).** Modification of HA with TBA salt is determined by the integration of the TBA methyl groups relative to the N-acetyl group of HA.

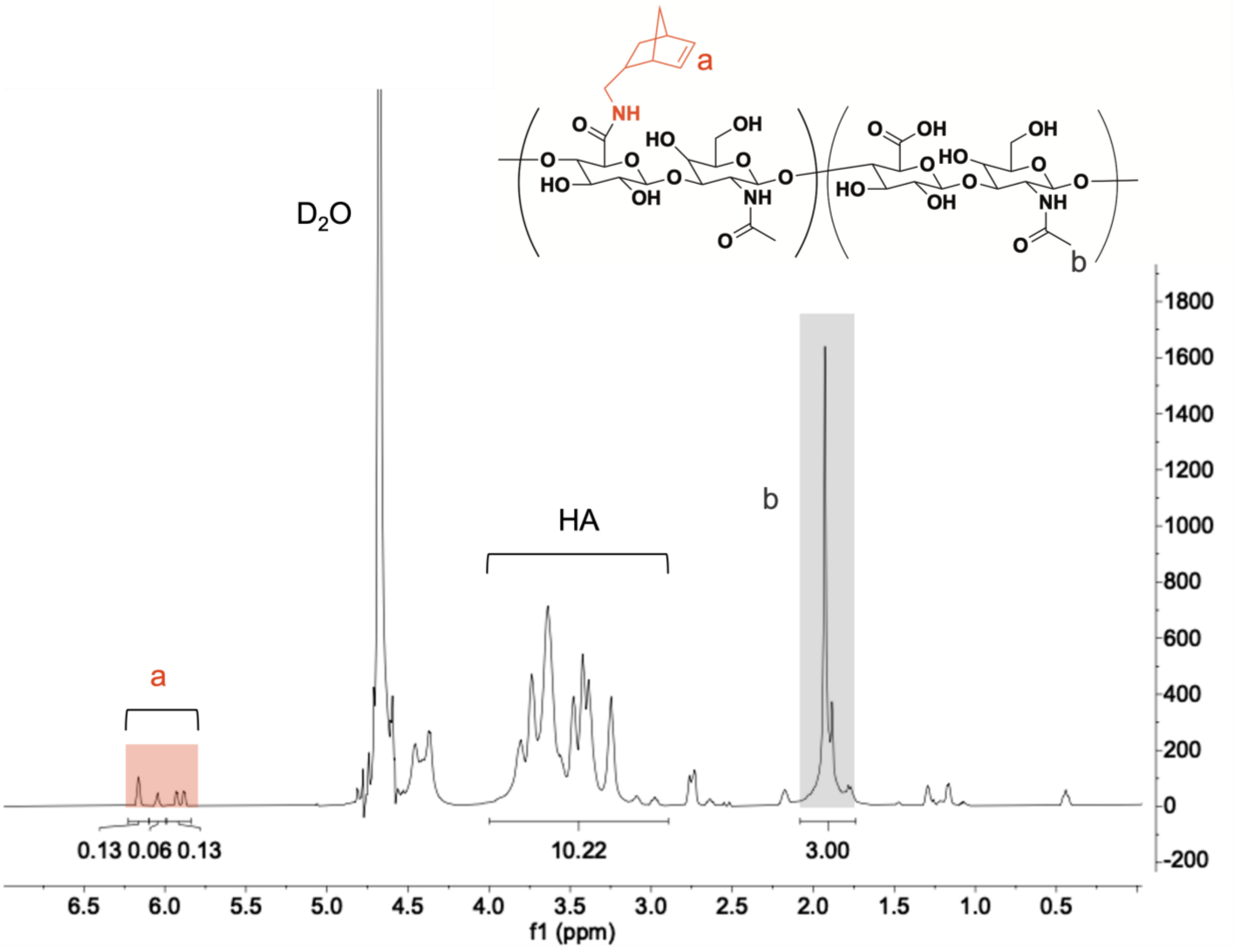


**Supplementary Figure 2: [1]H NMR spectrum of norbornene-modified hyaluronic acid (NorHA).** The degree of modification of HA with norbornenes was determined to be 16% as indicated by integration of the peaks at δ = 5.75, 6.05, and 6.17 (2H, 'a') relative to the N-acetyl on HA (3H, 'b').

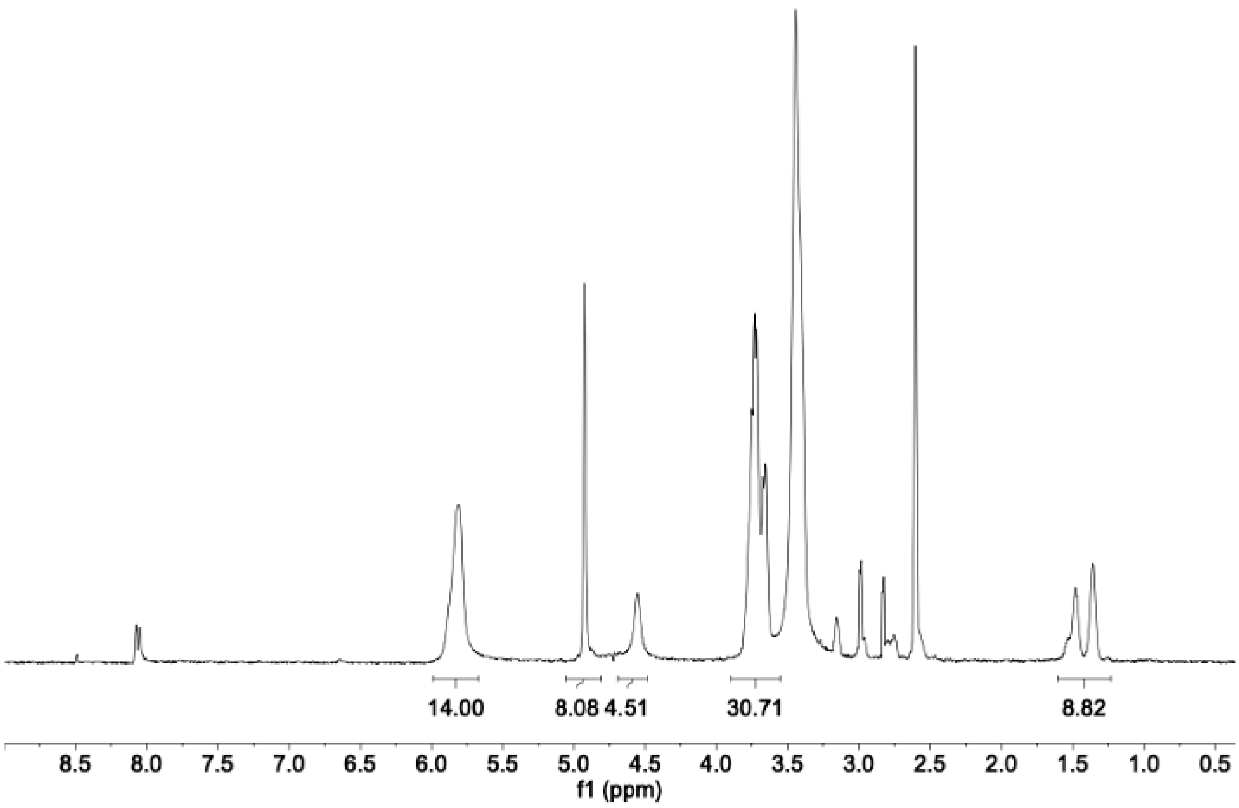


**Supplementary Figure 3: $^{1}$H NMR spectrum of β-cyclodextrin hexamethylene diamine (CD-HDA).** Modification of β-CD with HDA was determined to be 73% as indicated by integration of the peaks at δ = 1.14-1.6 ppm (12H).

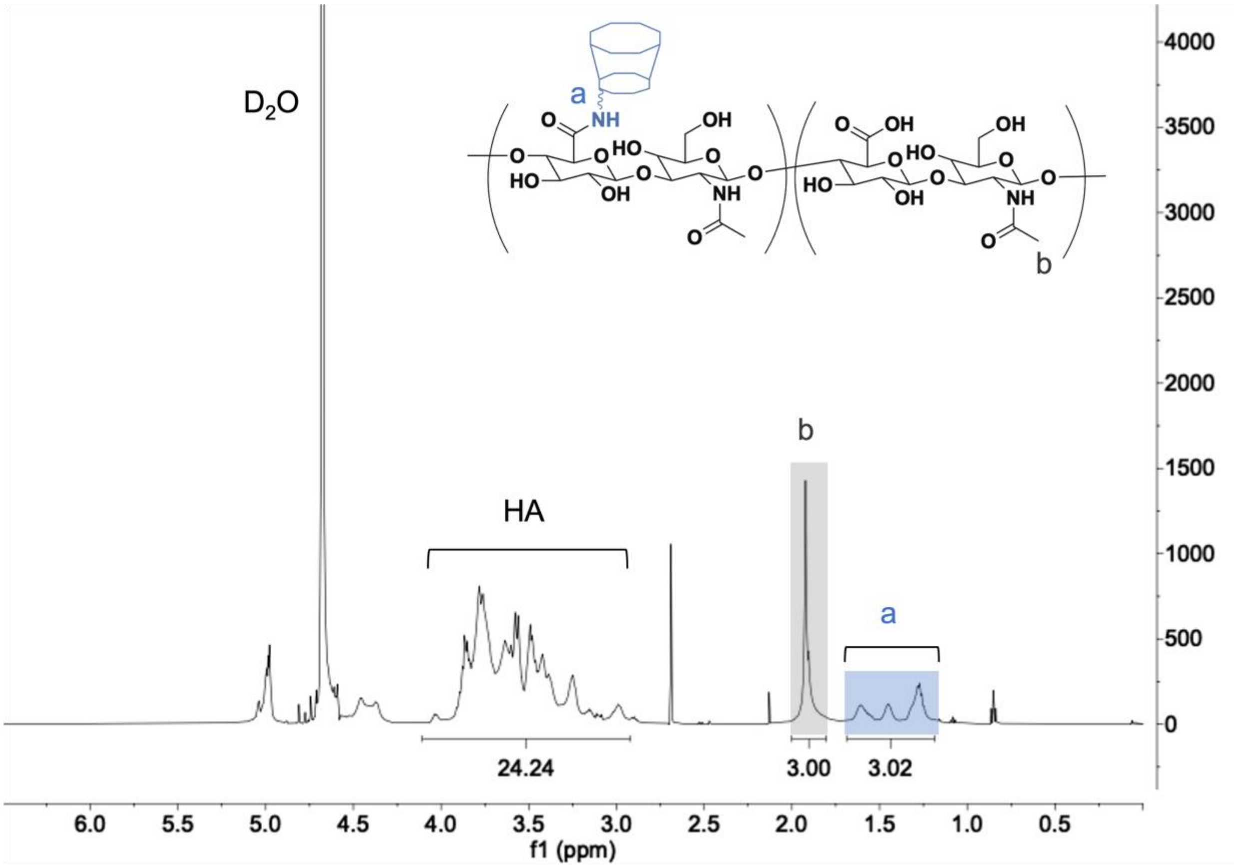


**Supplementary Figure 4: $^1$H NMR spectrum of β-cyclodextrin-modified hyaluronic acid (CD-HA)**. Modification of HA with pendant β-cyclodextrins (25%) was determined by integration of hexane linker peaks at δ = 1.23-1.68 ppm (12H, ‘a’) relative to the N-acetyl group of HA (3H, ‘b’).

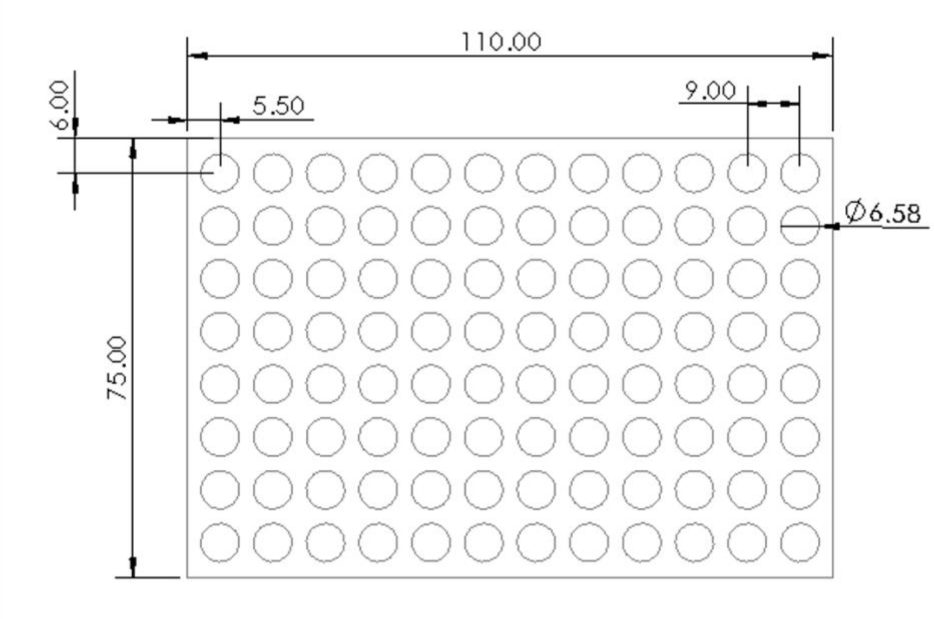


**Supplementary Figure 5: Dimensions used for laser cutting of the double-sided adhesive.** The dimensions shown were designed to match the bottom of the 96-well plate to which the adhesive was applied. All measurements are shown in millimeters.

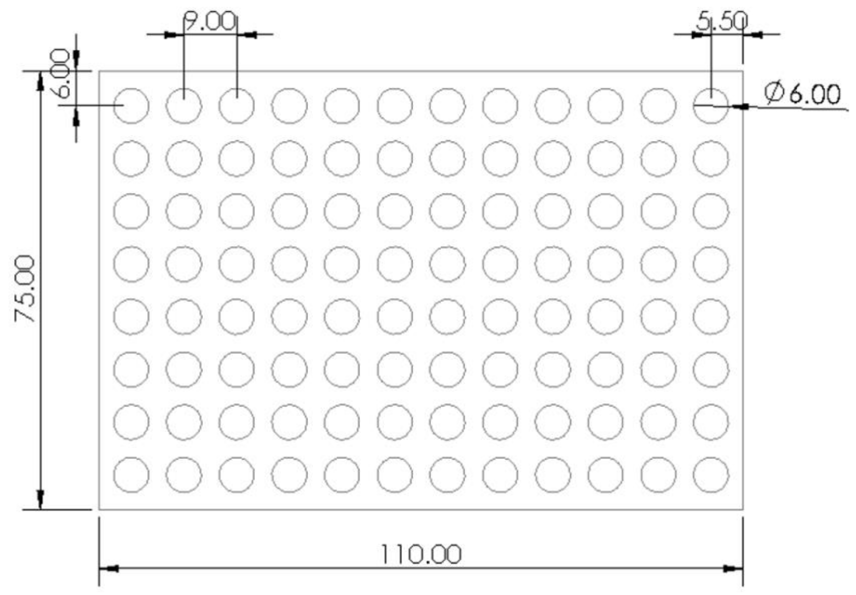


**Supplementary Figure 6: Dimensions used for laser cutting of the silicone spacer sheet.** The dimensions shown were designed to be about 0.5 mm less than the diameter of the well. All measurements are shown in millimeters.

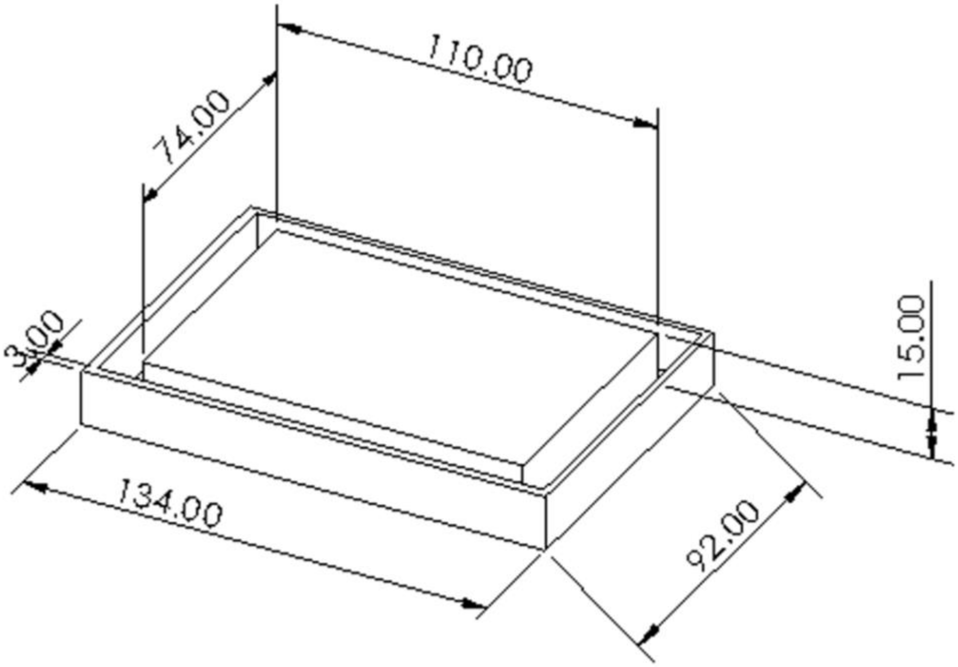


**Supplementary Figure 7: Dimensions of 3D-printed alignment piece used for 96-well hydrogel plate assembly.** This piece was 3D printed to assist with the alignment of the thiolated glass, silicone spacer, and bottomless 96-well plate for successful fabrication of the 96-well hydrogel array. All measurements are shown in millimeters.

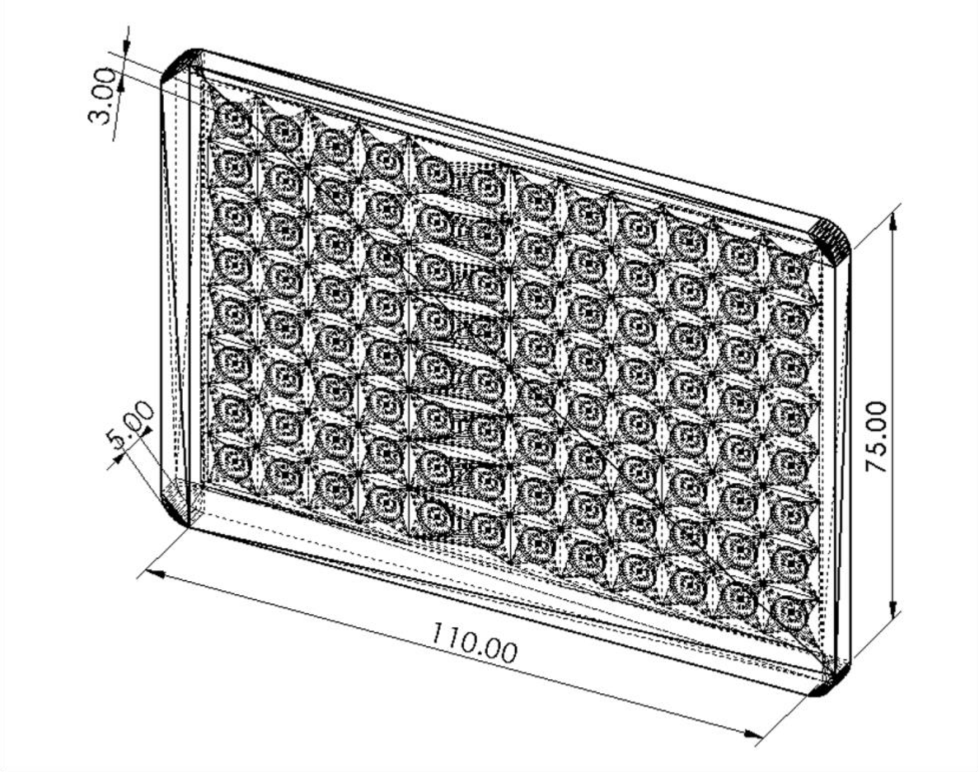


**Supplementary Figure 8: Dimensions of 3D-printed mold for fabrication of the PDMS spacer sheet.** This mold was 3D printed to assist in the fabrication of 1 mm thick PDMS sheets used to fabricate hydrogels for 3D culture within the 96-well array. All measurements are shown in millimeters.

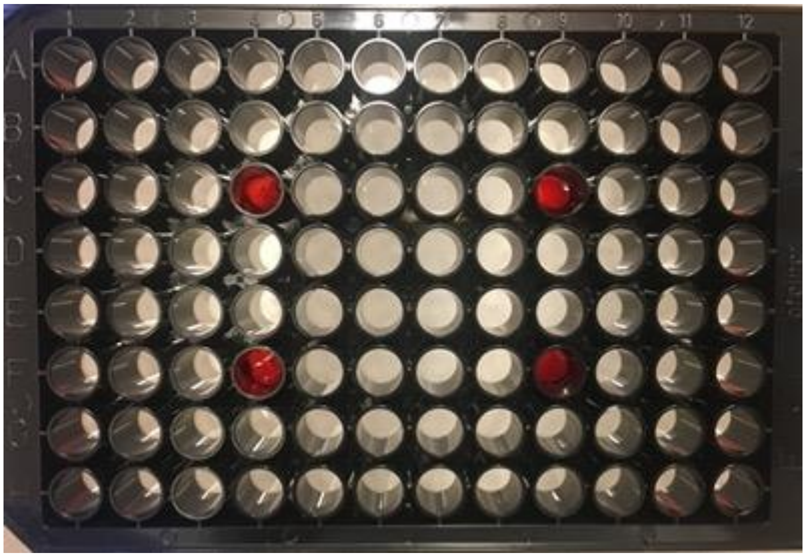

**Supplementary Figure 9: Leak test for 96-well plate hydrogel platform.** Four hydrogel wells in distinct regions of the 96-well plate were filled with red food coloring to visualize any potential liquid leakage between wells. This image was taken 24 hours after adding the food coloring, indicating that the liquid within the wells was well contained. Time lapse video of the leak test is provided in Supplementary Movie 1.
